# Optimal use of EEG recordings to target active brain areas with transcranial electrical stimulation

**DOI:** 10.1101/078758

**Authors:** Jacek P. Dmochowski, Laurent Koessler, Anthony M. Norcia, Marom Bikson, Lucas C. Parra

## Abstract

To demonstrate causal relationships between brain and behavior, investigators would like to guide brain stimulation using measurements of neural activity. Particularly promising in this context are electroencephalography (EEG) and transcranial electrical stimulation (TES), as they are linked by a reciprocity principle which, despite being known for decades, has not led to a formalism for relating EEG recordings to optimal stimulation parameters. Here we derive a closed-form expression for the TES configuration that optimally stimulates (i.e., targets) the sources of recorded EEG, without making assumptions about source location or distribution. We also derive a duality between TES targeting and EEG source localization, and demonstrate that in cases where source localization fails, so does the proposed targeting. Numerical simulations with realistic head models confirm these theoretical predictions and quantify the achieved stimulation in terms of focality and intensity. We show that constraining the stimulation currents automatically selects optimal montages that involve only a few (4-7) electrodes, with only incremental loss in performance. The proposed technique allows brain scientists and clinicians to rationally target the sources of observed EEG and thus overcomes a major obstacle to the realization of individualized or closed-loop brain stimulation.

## Introduction

The ability to systematically modify observed patterns of neural activity would be highly beneficial on at least two fronts: in basic neuroscience, mapping out the relationship between structure and function is facilitated by causal manipulations of brain activity. Moreover, techniques supporting target engagement provide novel strategies for treating psychiatric and neurological disorders marked by aberrant neural dynamics (Uhlhaas and Singer, 2006, 2012). An intriguing approach is to combine neuroimaging with brain stimulation (Bestmann and Feredoes, 2013; Bergmann et al., 2016; Siebner et al., 2009). The technical capability to perform integrated stimulation-recording of brain activity exists at a variety of scales: invasive microelectrode arrays (Maynard et al., 1997; Jimbo et al., 2003; Dostrovsky et al., 2000), deep brain stimulation (DBS) (Kent and Grill, 2013; Lempka and McIntyre, 2013; Rosin et al., 2011), depth electrodes (Rosenberg et al., 2009), cortical surface electrode arrays (Trebuchon et al., 2012), brain machine interfaces (Guggenmos et al., 2013), and non-invasive scalp electrode arrays that are commonly used in human neuroscience (Thut et al., 2005; Faria et al., 2012; Fernández-Corazza et al., 2016; Wagner et al., 2016). However, lacking is a general formalism for how to select stimulation parameters given observations of neural activity.

One particularly compelling combination is electroencephalography (EEG) with transcranial electrical stimulation (TES), mirror-symmetric processes related by the long-standing reciprocity principle introduced by Helmholtz (1853). Simply stated, the electrical path from a neural source to a (recording) electrode is equivalent to the electrical path from the (now stimulating) electrode to the location of the neural source (Rush and Driscoll, 1969). Intuition suggests that reciprocity should allow one to leverage the information carried by EEG signals to guide the parameters of the TES. Indeed, recent work has proposed *ad hoc* rules for distilling EEG measurements to TES configurations (“montages”) (Fernández-Corazza et al., 2016; Cancelli et al., 2016). However, these initial efforts have not realized the multi-dimensional nature of the reciprocity principle, and thus have failed to overcome the spatial blurring that results from naive implementations of reciprocal stimulation.

Here we develop a general formalism for combined EEG-TES, focusing on the problem of how to select the applied TES currents such that the source of an EEG activation is targeted by the stimulation. By formulating both EEG and TES as linear systems linked by a common transfer matrix, we derive a closed-form expression for the TES electrode configuration (“montage”) that generates an electric field most closely matched to the activation pattern. Importantly, we show that source localization of the targeted activation is not required, and that EEG sources may be stimulated using only their projections on the scalp. However, we also derive a duality between EEG localization and TES targeting, showing that the inherent limitations of localization are shared by targeting. In order to guarantee “safe” (i.e., current-limited) and feasible montages, we propose to constrain the *L*^1^ norm of the reciprocal TES solution, and provide a fast iterative scheme to achieve this.

In order to test the proposed approach, we conduct numerical simulations using a magnetic resonance imaging (MRI) based Boundary Element Model (BEM) of the human head. The simulations confirm the main theoretical prediction that in order to target the source of a recorded EEG pattern, the TES currents must be selected as the *spatially decorrelated* vector of measured EEG potentials. The duality between EEG and TES is also validated, and we present a high-noise scenario in which both EEG localization and TES targeting fail. We then demonstrate that when appropriately parametrized, the *L*^1^ constrained solution leads to simple montages yet provides an excellent tradeoff between the focality and intensity of stimulation. Finally, we also show that reciprocal stimulation accounts for varying source orientation, in that both radial and tangential sources are effectively targeted. In summary, we demonstrate that targeted stimulation of neural sources may be achieved by measuring neural activity at a surface array and using these measurements to design spatially patterned electrical stimulation. This approach has application to both human neuroscience and clinical interventions using neuromodulation.

## Results

TES delivers electric currents to the brain via an array of scalp electrodes, while EEG records voltages on the scalp generated by neural current sources in the brain. The goal of reciprocal TES is to select the stimulation currents on the scalp such that they reproduce the neural current sources in the brain. We provide the mathematical theory to optimally achieve this goal, while deferring proofs to the Methods. To test the theoretical predictions and estimate the performance of reciprocal TES in practice, we employ a BEM of the human head based on a tissue segmentation derived from MRI (see Methods for details). This “head model” allowed for the estimation of stimulation currents in the brain as well as simulation of voltage recordings due to neural currents.

### EEG lead field and TES forward model are symmetric

Consider an array of electrodes that is capable of both recording (neurally-generated) electric potentials and stimulating the brain with applied electrical currents. The recorded voltages, denoted by vector *V*, are a linear superposition of many neural current sources whose activity is represented by vector *J*:

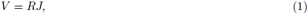

where *R* is the so-called “lead field” matrix (Sarvas, 1987; Hämäläinen and Ilmoniemi, 1994) that quantifies the voltages generated on the scalp by unit currents at various source locations in the brain. One example is given in Figure 1A, which shows a localized source of activity on the cortical surface. Note that the voltage recordings on the scalp are blurred due to volume conduction. The stimulation currents applied to the electrode array, denoted by vector *I*, generate an electric field *E* inside the brain:

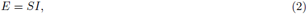

where matrix *S* is the “forward model” (Dmochowski et al., 2011) that quantifies the electric field generated in the brain for a unit current applied to each of the stimulation electrodes. In this multiple electrode context, reciprocity leads to a symmetry relationship among *R* and *S*:

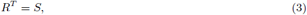

where ^*T*^ denotes matrix transposition. This formulation of reciprocity is novel in that it describes the relationship between *multiple* neural sources and *multiple* electrode pairs. Reciprocity for individual sources and a single pair of recording electrodes in a non-uniform medium such as the brain has been known for decades (Rush and Driscoll, 1969), and linear superposition of multiple sources has been previously leveraged for current flow modeling (Hallez et al., 2007; Huang et al., 2015; Wagner et al., 2016), but a compact formulation as in Equation (3) was lacking. We provide a derivation for this multidimensional reciprocity in the Methods. In the next section, we exploit multi-dimensional reciprocity to, for the first time, selectively target active neural sources with appropriately tuned stimulation currents.

**Figure 1.**
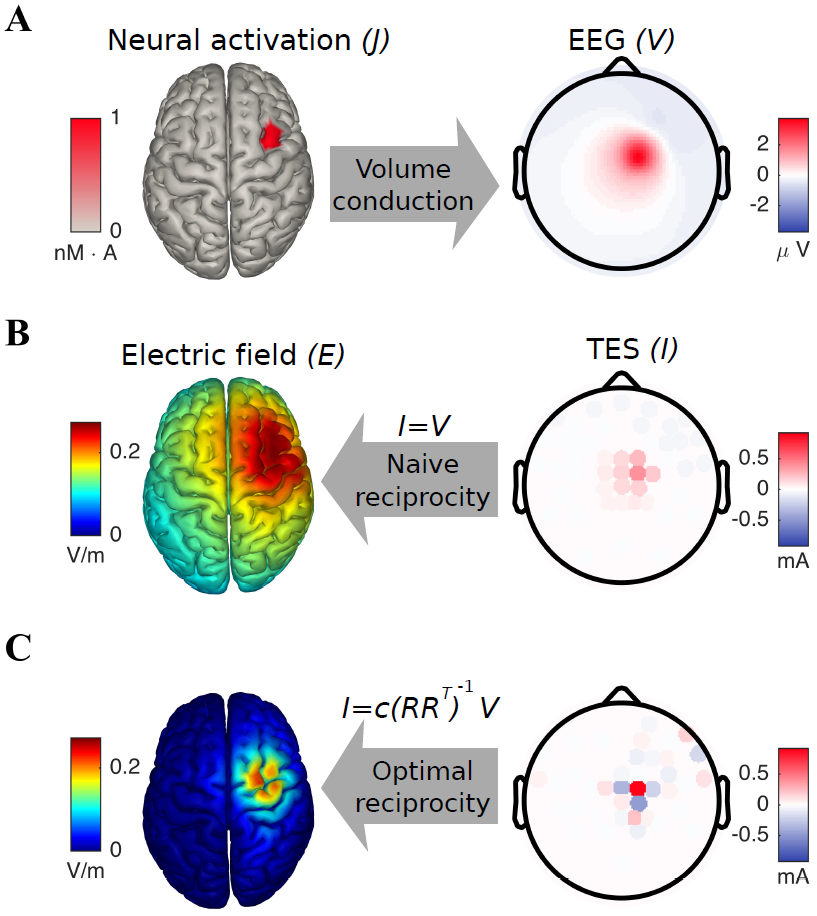
Reciprocal stimulation produces an electric field focused on the site of neural activation. (**A**) Focal neural activation of the right frontocentral cortex produces a radially-symmetric pattern of electric potentials on the scalp. (**B**) By patterning the stimulation currents according to the observed scalp activity (i.e., *I* ∝ *V*), “naive” reciprocity generates a diffuse electric field that is strong at the site of activation but also over expansive regions of cortex. (**C**) Applying TES in proportion to the spatially decorrelated EEG (i.e., *I = c(RR^T^)^−1^ V)* yields focal stimulation at the neural activation. Note that the injected reciprocal currents are both positive (“anodal”) and negative (“cathodal”) over the scalp regions marked by positive EEG potentials.

### Reciprocal TES inverts the spatial blurring of EEG potentials

To modulate the neural activity underlying the EEG, we propose to recreate with TES an electric field that matches the neural source distribution. In mathematical terms, an ideal outcome is thus *E = cJ*, where *c* is a constant that relates the magnitude of neural activation (measured in A · m) to the strength of the desired electric field (measured in V/m). The selection of stimulation currents to achieve this goal can be formulated as a convex optimization problem:

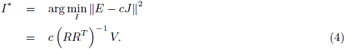

The result of (4) states that to modulate the sources of an observed EEG pattern *V*, one should apply TES currents according to *c (RR^T^)^−1^ V*, where the matrix inverse compensates for the spatial mixing due to lead fields *R*. This solution is in contrast to a “naive” reciprocity approach, which simply applies currents with the same spatial pattern as the recorded voltage distribution: *I^*^ ∝ V* (Cancelli et al., 2016) and does not decorrelate the recorded voltages. Note that the ability of optimal reciprocity (4) to account for volume conduction is predicated on the availability of the lead field matrix *R*, which conveys the set of *possible* source locations (e.g., all cortical locations).

To compare the results of optimal reciprocity with the naive approach, we simulated activation of a patch of cortical tissue (bounded by a sphere of 1 cm radius) in the right frontal cortex (Fig 1A, left). Each source was oriented perpendicular to the cortical sheet, following the notion that pyramidal neurons are the primary source of the EEG (Nunez and Srinivasan, 2006). The resulting scalp potentials were marked by a positivity at right frontocentral electrodes and a diffuse negativity at surrounding sites (Fig 1A, right). This EEG pattern was then used as an input into two forms of TES aimed at stimulating the activation region: the naive reciprocity approach (*I^*^ ∝ V*) and optimal reciprocity according to (4). The electric field resulting from the naive approach (Fig 1B, left) had a large magnitude of 0.25 V/m in the neural activation region; however, the field pattern was excessively diffuse. The focality of the electric field, quantified as the radius at which field magnitude drops by half, was 8.2 cm (Fig 1B, left). Optimal reciprocity resulted in a focused electric field that still strongly activated the target region (0.16 V/m), while exhibiting a more compact stimulation radius of 4.0 cm (Fig 1C, left). The resulting field did not exactly match the original neural source distribution because the scalp potentials were only measured at a limited number of locations (i.e., 64). In theory, if voltages could be measured noise-free and with as many electrodes as neural sources, one could exactly recreate the distribution.

Note that the reciprocal TES montage consisted of both positive and negative stimulation currents in the scalp region marked by positive EEG (Fig 1C, right). We also mention that the result of (4) corresponds to “anodal” stimulation: that is, direct currents I will depolarize neurons oriented perpendicular to the cortical surface. If one instead seeks to hyperpolarize such cells (“cathodal” stimulation), the stimulation currents take the form *I^*^ = −c (RR^T^)^−1^ V*.

### EEG localization and TES targeting are dual problems

Note that the neural source distribution *J* does *not* enter the expression for the reciprocal currents *I^*^*: this variable is “absorbed” by the lead field R in the derivation of (4)-see the Methods. This means that in order to reproduce the activity pattern, one need not know the locations of the active sources in the brain, but only their voltage measurements on the scalp. It may thus appear that reciprocity has solved the ill-posed inverse problem inherent to encephalography (Pascual-Marqui, 1999). However, as we show in the Methods section, source localization and reciprocal targeting are actually dual mathematical problems. Specifically, the optimal electric field E* achieved with reciprocal targeting following (4) is proportional to the conventional minimum-norm estimate of the neural source distribution *J^*^*:

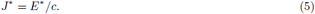

Importantly, (5) implies that in instances where the minimum-norm source estimate *J^*^* is inaccurate, the electric field *E^*^* generated by reciprocal stimulation will also be misguided.

To examine the duality between source localization and reciprocal targeting, we simulated bilateral activation of the parietal cortex (Fig 2A) under two distinct noise conditions: spatially white noise was added to the electrodes with a signal-to-noise ratio (SNR) of either 100 or 1. In the low-noise case, the topography of the EEG clearly showed a radially symmetric pattern centered over centroparietal electrodes (Fig 2B). The TES montage that reciprocated this pattern consisted of a central anode surrounded by four cathodes (Fig 2C), producing an electric field with an intensity of 0.12 V/m at the activated region and a focality of 3.3 cm (Fig 2D). We then computed the minimum norm source estimate of the EEG topography. Confirming the theoretical prediction (Eq. 5), this estimate was found to be perfectly correlated with the electric field generated by reciprocal TES (*r* = 1, Fig 2E). In the high-noise case, the EEG topography exhibited distortion (Fig 2F), leading to a reciprocal TES montage that erroneously recruited bilateral frontal electrodes (Fig 2G). The electric field produced by this montage “missed” the site of activation and was distributed along the midline (Fig 2H). Again confirming the theoretical prediction, the minimum-norm estimate of the neural source distribution underlying the distorted EEG pattern was found to be perfectly correlated with this electric field (*r* = 1, Fig 2I).

**Figure 2.**
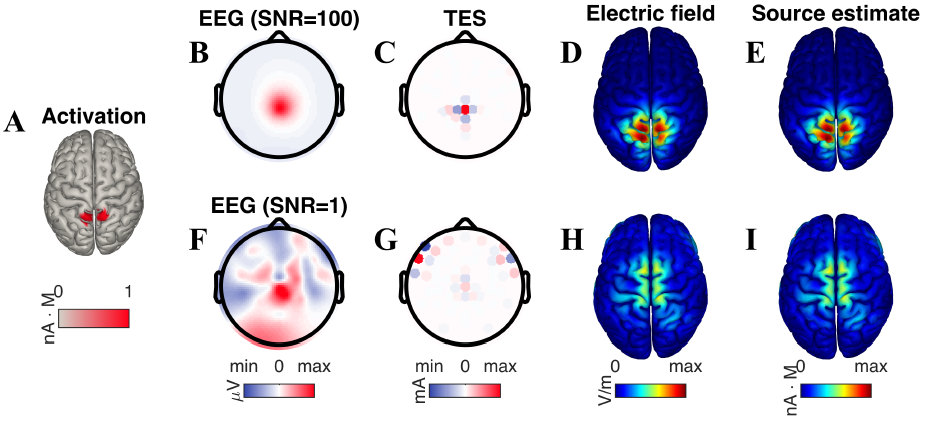
Localization of EEG is equivalent to targeting in TES. (**A**) Bilateral activation of the superior parietal lobule. (**B**) The observed EEG pattern, simulated here for a high signal-to-noise ratio (SNR) of 100, shows a radially-symmetric topography focused over centroparietal electrodes. (**C**) The TES montage that targets the source of this EEG is composed of a center anode and surrounding cathodes, producing an electric field (**D**) concentrated at the source of the parietal activation. Moreover, this electric field is perfectly correlated with the minimum-norm estimate (**E**) of the EEG source distribution. (**F**) An increase in the noise level (SNR=1) leads to a distorted EEG topography, which then results in a reciprocal TES montage (**G**) that erroneously utilizes lateral frontal electrodes. The resulting electric field (**H**) is no longer focused on the site of neural activation. Correspondingly, the estimate of the EEG source (**I**) is also mismatched with the actual neural activation.

### Constraining reciprocal TES leads to safe and feasible stimulation

The magnitudes of *I*^*^, as computed by Eq. (4), may be less or greater than those desired in practice. For example, the presently accepted safety practice for TES is to deliver no more than 2 mA to the head (Brunoni et al., 2012; Bikson et al., 2016). Denoting the limit on current delivered by *I_max_* (e.g. 2 mA), safe stimulation corresponds to a constraint on the sum of absolute currents:

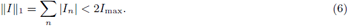

Eq. (6) specifies a constraint on the “*L*^1^ norm” of stimulation currents *I*. In cases where the unconstrained solution from Eq. (4) violates the inequality, we must somehow adjust *I^*^* to adhere to this safety constraint. The simplest way of achieving this is through a uniform scaling of the elements of *I^*^*:

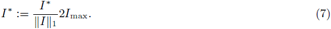

However, it is important to note that scaling *I^*^* via (7) does *not* minimize the mean squared error between *E* and *cJ* subject to the constraint (6). Therefore, we propose the following constrained optimization problem:

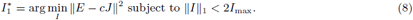

Unlike the unconstrained case (4), the optimization problem (8) does not have a closed-form solution. However, a number of algorithms have been proposed in order to numerically solve such *L*^1^-constrained least-squares problems (Tibshirani, 1996). In the Methods, we propose an iterative scheme that converts the (non-differentiable) *L*^1^ constraint to a set of linear constraints that may be iteratively solved using standard numerical packages. The solution to (8) represents the pattern of stimulation currents that best recreates the neural activation while maintaining safe current levels.

To evaluate the effect of constraining the *L*^1^ norm of the reciprocal TES solution on the achieved stimulation, we simulated activation of four distinct cortical regions: the superior temporal gyrus (STG, Fig 3A), the dorsolateral prefrontal cortex (DLPFC, Fig 3B), the superior parietal lobule (SPL, Fig 3C), and visual area V5 (also known as the middle temporal (MT) visual area, Fig 3D). In the evaluation, we examined the role of parameter *c*, which relates the strength of the desired electric field *E* to the magnitude of the neural activation *J*. For unconstrained reciprocity (4), the value of *c* is inconsequential-in practice, the currents will be scaled to the desired level using Eq. (7). However, for *L*^1^-constrained reciprocity (8), the value of *c* will determine the distribution of the optimal stimulation currents, and consequently, the shape of the electric field. Given that the neural sources of the EEG have intensities in the order of 1 nm. A (Nunez and Srinivasan, 2006), and that the electric fields produced by TES are in the order of 0.5 V/m (Opitz et al., 2016), the parameter should be selected to be large (i.e., *c* ranged from 10^7^ to 10^13^). Larger values of c led to higher intensity but lower focality of stimulation at the target (Fig 3A-D, circle markers). Importantly, the focality-intensity curve exhibited a sharp “knee”, indicating that by carefully selecting *c*, stimulation intensity can be greatly increased while only suffering a minimal reduction in focality. For the example activations here, the knee point occurred at *c* = 10^10^. In addition to increasing stimulation intensity, larger values of c also produced montages utilizing less active electrodes (represented by color of markers). At the knee point, *L*^1^-constrained montages recruited between 4 and 7 electrodes. The electric field achieved by unconstrained reciprocity emphasized electric field focality at the expense of intensity (Fig 3A-D, diamond marker).

**Figure 3.**
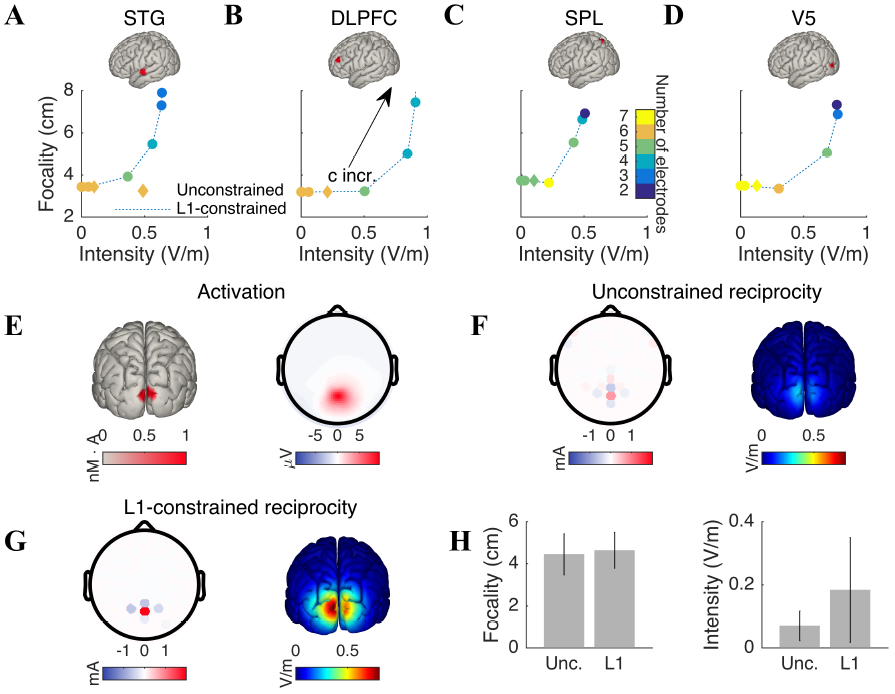
*L*^1^-constrained reciprocity produces safe, feasible, and intense stimulation. Reciprocal TES at increasing values of parameter c was performed on the EEG generated by activation of the (**A**) superior temporal gyrus (STG), (**B**) the dorsolateral prefrontal cortex (DLPFC), (**C**) superior parietal lobule (SPL) and (**D**) visual area V5 (also known as the middle temporal visual area). Increasing c led to higher intensity and reduced focality at the activation. However, the focality-intensity curve exhibited a sharp “knee” point at *c* = 10^10^, meaning that by choosing c appropriately, stimulation intensity can be increased without reducing focality. Increasing c also led to a reduction in the number of electrodes recruited by the optimal montage (denoted by color of markers). Unconstrained reciprocity produced stimulation that emphasized focality (diamond marker). (**E**) Activation of primary visual cortex and the associated EEG pattern. (**F**) Unconstrained reciprocity distributed the applied current over approximately 8 electrodes. This led to a concentration of the electric field at the occipital target (focality of 3.3 cm), with a peak electric field intensity of 0.18 V/m. (**G**) By constraining the *L*^1^-norm of the reciprocal TES solution with *c* = 10^10^, the applied currents are contained to 5 electrodes, yielding to a three-fold increase in the intensity of the stimulation at the activated region (0.53 V/m), while only sacrificing 4 mm in focality (3.7 cm). (**H**) Comparing unconstrained and *L*^1^-constrained reciprocity across all cortical sources demonstrates that *L*^1^-constrained reciprocity achieves an average increase in field intensity of 163%, while only sacrificing 4% in focality (error bars represent standard deviations across 15,002 sources).

### Intensity can be increased without reducing focality

To test whether choosing a value of *c* near the knee point brings about the predicted increase in intensity, we performed *L*-constrained reciprocity with *c* = 10^10^ on activations from all sources in the head model (see Methods for details). For each activation, we computed both the optimal (unconstrained) and *L*^1^-constrained reciprocal TES montages. An example activation of the primary visual cortex is depicted in Fig 3E. In this case, unconstrained reciprocity led to the stimulation currents being distributed over approximately 8 electrodes, producing an electric field with an intensity of 0.18 V/m at the activation and a focality of 3.3 cm (Fig 3F). The *L*^1^-constrained montage was limited to 5 active electrodes, and produced a three-fold increase in the mean electric field intensity at the activated region (Fig 3G, 0.53 V/m), while only suffering a slight reduction in focality (i.e., 3.7 cm). Comparing the focality and intensity of the electric fields produced by both unconstrained and *L*^1^-constrained reciprocity across all activations confirmed that the *L*^1^-constrained solution provides an excellent tradeoff between focality and intensity (unconstrained reciprocity: 0.07 ± 0.05 V/m, 4.5 ± 1.0 cm; *L*^1^-constrained: 0.18 ± 0.17 V/m, 4.6 ± 0.8 cm; Fig 3H). The difference between unconstrained and *L*^1^ constrained was found to be statistically significant in both focality (*p* = 0 to numerical precision, *N* = 15002, Wilcoxon signed rank test), and intensity (*p* = 0).

### Performance of reciprocal TES is best at dorsal targets

In order to quantify the performance of reciprocal TES as a function of target location, we plotted the focality and intensity achieved by *L*^1^-constrained reciprocity (see Fig 3H) on cortex. The focality of stimulation ranged from 2.9 to 7.7 cm and exhibited optimal values over broad regions of the dorsal cortical surface (Fig 4A). Electric fields were the least focal for sources on the ventral temporal surface. Unlike focality, stimulation intensity showed an idiosyncratic spatial distribution, with discrete “hotspots” appearing on the dorsal surface of all four lobes (Fig 4B). The peak electric field magnitude of reciprocal TES was found to be 0.71 V/m, with intensity dropping off sharply along the ventral surface. As expected, a strong and significant negative correlation was observed between focality and intensity (*r* = −0.56, *p* = 0 to numerical precision, *N* = 15, 002).

**Figure 4.**
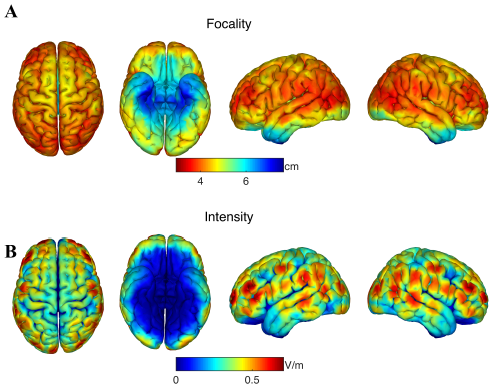
The performance of reciprocal stimulation is best at dorsal targets. (**A**) The focality of **L*^1^*-constrained reciprocal TES, as measured by the radius bounding half of the total electric field, shown as a function of the location of neural activation. Focality peaks at 2.9 cm and exhibits low variability along the dorsal surface of cortex. On the other hand, targeting activations originating on the ventral surface leads to significantly decreased focality (up to 8 cm). (**B**) Same as A but now for the intensity of the electric field at the target. The magnitude of stimulation exhibits discrete “hotspots” along the dorsal surface, peaking at 0.71 V/m. As with focality, intensity is limited when targeting activations of the ventral cortex. Locations receiving focal (diffuse) stimulation are stimulated with high (low) intensity (correlation between focality and intensity at target: *r = −0.56, p = 0, N = 15, 002*).

### Reciprocity accounts for source orientation

A leading hypothesis in TES research (at least for direct current stimulation) is that polarization of neurons is dependent on the direction of the applied electric field relative to the affected cells (Rahman et al., 2013). For example, maximal stimulation of pyramidal cells is achieved when the electric field is aligned with the somatodendritic axis (Bikson et al., 2004; Radman et al., 2009) We therefore sought to determine whether the electric fields generated by reciprocal TES are matched to not only the location but also the orientation of the activated sources. To test this, we simulated focal activation of a gyral patch of motor cortex (Fig 5A) with two orientations: (i) normal to the local cortical surface and (ii) tangential to the local surface (i.e., left to right). We performed *L*^1^-constrained reciprocal TES for both cases and determined the electric field vector at the activated region. Radial activation led to a largely monopolar EEG pattern concentrated at the left frontocentral electrodes (Fig 5B). Reciprocal stimulation for this scalp topography consisted of a dominant central anode and three surrounding cathodes (Figure 5C), and produced an electric field with a strong radial component (0.24 V/m, Fig 5D, activation region indicated by white circle). The field’s tangential component had a weaker intensity (0.09 V/m, Fig 5E), meaning that stimulation would have produced significantly more polarization of normally oriented tissue, thus matching the source orientation. Tangential activation resulted in an antisymmetric dipolar pattern focused over central electrodes (Fig 5F). In this case, the dominant anode was more left lateralized, with an idiosyncratic pattern of cathodes (Fig 5G). Importantly, the tangential component of the field was 0.30 V/m (Fig 5H), while the field strength in the radial direction was only 0.12 V/m (Fig 5I). Thus, for both radial and tangential activations, the dominant orientation of the reciprocal electric field matched the direction of the activated tissue.

**Figure 5.**
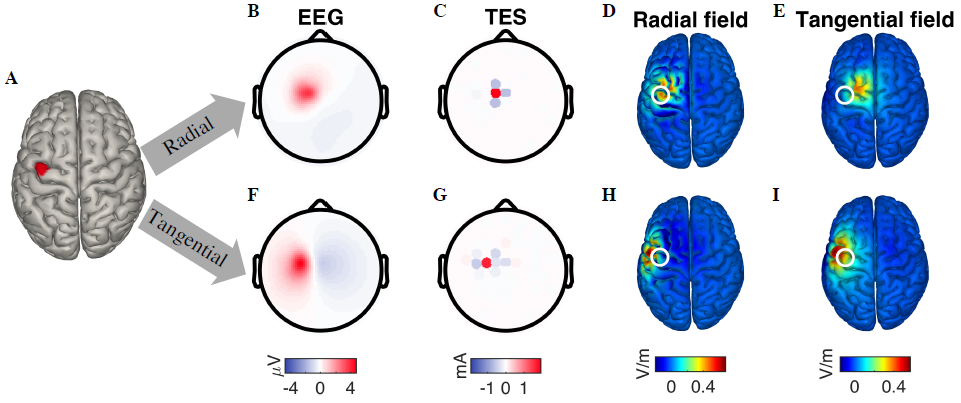
Reciprocal TES accounts for varying source orientation. (**A**) A source in the left motor cortex was simulated with both radial and tangential orientations. (**B**) Radial activation led to a monopolar EEG pattern over frontocentral electrodes. (**C**) The reciprocal TES montage for this scalp pattern consisted of a central anode with three surrounding cathodes. (**D**) The resulting electric field was marked by a strong radial component (0.24 V/m) and a significantly weaker (**E**) tangential component (0.09 V/m). (**F**) Tangential activation resulted in a broad antisymmetric pattern of scalp potentials. (**G**) The TES montage targeting this EEG pattern consisted of a more left-lateralized anode and multiple weak cathodes. (**H**) This montage produced a weak radial electric field component (0.12 V/m) relative to the (**I**) tangential direction of the electric field (0.30 V/m). Thus, for both cases, reciprocal TES produced an electric field whose dominant direction matched the orientation of the activated source.

## Discussion

### The duality of TES and EEG

Here we show how measured EEG potentials may be translated into optimal TES montages that generate electric fields focused over the areas of neural activity. To target the sources of observed EEG, we describe a pattern of TES currents matching the *spatially decorrelated* scalp potentials. In contrast, the simplistic strategy of placing the anodes over positive potentials and the cathodes over negative potentials results in drastically suboptimal stimulation (see Fig 1). Importantly, the determination of the reciprocal TES montage does not require source localizing the measured EEG. However, this does *not* imply that reciprocity has solved the ill-posed inverse problem of EEG. Rather, it was shown that EEG source localization and TES targeting are in fact two sides of the same optimization problem. Thus, when trying to reciprocate an EEG topography that fails to yield a meaningful source estimate via minimum-norm localization (Sarvas, 1987; Hämäläinen and Ilmoniemi, 1994), the resulting TES targeting “misses” the neural activation (see Fig 2). Reciprocal TES allows one to stimulate EEG sources using measurements of their activity on the scalp, but with the caveat that the limitations of the targeting mirror those of source localization. Nonetheless, the methods proposed here are numerically optimized given these biophysical constraints. Moreover, our approach prescribes a clear implementation that is applicable to any EEG, while simplifying hardware by minimizing TES electrodes. The approach proposed here is not limited to transcranial stimulation. It applies identically to the case of deep brain stimulation that is guided by electrical recordings of neural activity inside the brain.

### The need for a head model

In order to guide TES using EEG, computation of the lead-field matrix R, or equivently, the forward model S, is required. These matrices specify the possible locations of the EEG sources that may be targeted by stimulation. Despite the availability of freely-available tools for constructing forward models (Thielscher et al., 2015; Jung et al., 2013; Truong et al., 2014), their widespread adoption for designing TES interventions has not transpired. One potential reason for this is the high cost of acquiring structural MRI scans for study participants. In this case, one can opt to use a template head model, such as the one used here (Mazziotta et al., 1995) or the dense array version (Huang et al., 2015). An important question that has not been answered here is how much the performance of reciprocal TES degrades when applying a template model to individualized EEG recordings. It should also be noted that due to reciprocity, lead fields routinely computed for EEG (Tadel et al., 2011; Gramfort et al., 2014) may be equivalently used for TES targeting. One caveat here is that while EEG sources are typically restricted to the grey matter, the target of a TES study may be subcortical. In such cases, however, the EEG is unlikely to be used to inform TES, as the contribution of subcortical areas to the EEG is presumed to be small.

### Benefits of *L*^1^-constrained reciprocity

The stimulation currents computed by unconstrained reciprocity (4) may differ in scale from those desired in practice (e.g. 2 mA total). Here we showed that imposing an inequality constraint on the total current led to reciprocal TES montages that adhered to this desired scaling while still focally stimulating the neural sources (see Fig 3). In fact, *L*^1^-constrained reciprocity produced stimulation that would likely be favored in practice over its unconstrained counterpart: when properly selecting the parameter *c*, the intensity more than doubled while sacrificing only a fractional amount of focality (see Fig 3). The optimal value of *c* (which models the ratio of intensities between TES electric fields and EEG sources) found here (10^10^) is roughly consistent with present estimates for EEG source intensities (1 nm · A) and TES electric fields (0.5 V/m). In addition to increased intensity, *L*^1^-constrained montages are also more feasible, as they recruit a limited number of electrodes (see Fig 3). This is a natural consequence of imposing *L*^1^ constraints, which are well-known to yield sparse solutions to least squares problems (Tibshirani, 1996).

### Comparison with recent work

The reciprocal TES approach described here leverages EEG recordings. It differs therefore fundamentally from other forms of TES targeting that are based purely on anatomical information (Dmochowski et al., 2011). A previous attempt to leverage EEG for targeting (Cancelli et al., 2016) is based on the intuition that the injected currents should match the recorded voltages (*I ∝ V*), which we referred to here as the “naive” reciprocity approach (Figure 1B) as it does not recognize the importance of inverting the blurring introduced by volume conduction. Fernández-Corazza et al. (2016) suggest the use of the traditional reciprocity principle, but fail to recognize the multi-dimensional reciprocity relationship (3). As a result, the EEG and TES problems could not be mathematically synthesized into the least-squares optimization problem developed here (4). Thus, the solutions of Fernández-Corazza et al. (2016) are limited to heuristics relying on individual electrode pairs. Consequently, neither of these two previous efforts were able to focus the electric field onto the site of neural activation. In contrast, the approach presented here is optimal in reproducing the neural source distribution in a least-squares sense.

### Practical implementation

The EEG is almost always acquired over multiple electrodes and time points, and is thus commonly represented as a space-time data matrix. Reciprocal TES takes as an input a vector of scalp potentials, meaning that the EEG matrix must first be distilled into a time-independent vector. This can be accomplished in several ways. The simplest is to select a time point and use a temporal slice of the data as the input: in this case, the stimulation will be focused on the regions whose activation is strongest during the selected time point. To lessen the sensitivity of the scheme to the particular time point chosen, an alternative approach is to temporally average the EEG across some epoch of the data (e.g. the length of an experimental trial)-in this case, reciprocal TES will target sources most strongly expressed over the epoch duration. Yet another possibility, and one that is most principled, is to decompose the data into spatial components via a technique such as principal components analysis (PCA) (Parra et al., 2005), independent components analysis (ICA) (Delorme and Makeig, 2004), or reliable components analysis (RCA) (Dmochowski et al., 2012). In this case, one can inspect the topographies of the various components in order to select the one that is to be targeted with reciprocal stimulation. It is important to note that when reciprocating a component of the EEG, one should use the forward projection of the weights and *not* the spatial filter weights themselves (Parra et al., 2005; Haufe et al., 2014).

### Physical limits on performance

Given constraints on the number of electrodes (i.e., 64) and the total current delivered (i.e., 2 mA), the maximum achievable focality of reciprocal TES was here found to be approximately 3 cm. As defined here, this means that half of the total electric field was confined to a sphere of 3 cm radius. Optimal focality was broadly achieved across the dorsal surface of cortex, and dropped off steeply on the ventral surface (i.e., a focality of almost 8 cm). The strongest electric field intensity was found to be 0.7 V/m, with these maximal values located at discrete “hotspots” along dorsal patches of the cortical surface. These hotspots likely represent locations with a favorable electrical channel from scalp to cortex (and reciprocally, from cortex to scalp). For both focality and intensity, reciprocating sources located along the ventral surface was found to be challenging, with simultaneously low focality and intensity produced by reciprocal TES to those regions. The inclusion of low-lying electrodes (for example, on the neck) that provide more coverage of ventral surfaces may increase our ability to effectively stimulate ventral brain regions.

### Implication for closed-loop TES

Closed loop brain stimulation, during which neural recording and stimulation are performed simultaneously or in tandem, has already been shown to be effective in reducing pathophysiological patterns in Parkinson’s Disease (Rosin et al., 2011) and blocking epileptic seizures (Berényi et al., 2012; Osorio et al., 2001). In the context of non-invasive techniques, transcranial alternating current stimulation (TACS) has been reported to entrain oscillatory EEG rhythms (for example, alpha oscillations) (Antal and Paulus, 2013), including in an open-loop design taking into account the individual oscillation frequency of the subjects (Zaehle et al., 2010). Unlike these prior techniques, the reciprocal approach developed here uses the spatial (i.e., as opposed to temporal) statistics of neural activity to guide stimulation. As this spatial information is orthogonal to temporal dynamics, our approach may be performed on both oscillatory and evoked activity. Moreover, it could also be combined with existing approaches that tune the temporal frequency of the applied stimulation. While we appear to have a solid understanding of electrical fields generated in the brain during stimulation (Opitz et al., 2016), it may be argued that we know less about how these electric fields interact with neuronal activity. Closed-loop stimulation efforts will be enhanced by mechanistic knowledge of the micro-and meso-scale interactions between biological activity and the modulating electric fields. In the context of EEG-TES, this entails developing a functional link between the parameters of the stimulation and the resulting changes to the EEG, which is perhaps mediated by the brain state at stimulation onset (Silvanto et al., 2008).

## Methods

Here we provide the proofs of the theoretical findings presented in the Results. In particular, by leveraging linear superposition of electric potentials, we derive a vectorial form of the well-known reciprocity between electrical stimulation and recording (Rush and Driscoll, 1969), relating here not a single current source in the volume with a pair of electrodes, but rather an array of electrodes with multiple current sources. We then employ this multidimensional reciprocity to solve for the array of scalp currents that best targets the source of an observed EEG pattern.

### Theorem 1.

Multi-dimensional reciprocity.

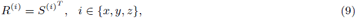

where *R^(i)^* and *S^(i)^* denote the *i*th Cartesian components of the lead-field matrix and forward model, respectively.

*Proof.* (Theorem 1) The fundamental reciprocity relation, when written for the case of a single source in the brain and a single electrode pair on the scalp is given by (Rush and Driscoll, 1969):

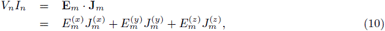

where *V_n_* is the voltage at scalp location *n* (relative to a reference electrode) due to a current source **J**_*m*_ at brain location *m* (bold font indicates that this is a three-dimensional vector in Cartesian space *{x,y,z}*), and reciprocally, **E**_*m*_ is the electric field vector generated at brain location *m* when applying a current of *I_n_* to scalp location *n* (and hence a current of −*I_n_* at the reference electrode). Moreover, we have explicitly written out the *x-, y-*, and *z-* components of the electric field as 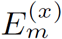, 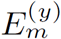, and 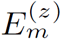, respectively, with analogous definitions holding for the current source vector **J**_*m*_.

We seek to extend this result to the case of multiple sources in the brain (indexed by *m*) and multiple electrode pairs on the scalp (indexed by *n*). The fundamental observation is that under quasi-static conditions, the electric potentials and fields from multiple current sources are additive (Jackson, 1999). Invoking this superposition principle on the component voltages *V_nm_* generated at electrode n by multiple brain sources **J**_*m*_, as given by (10), yields:

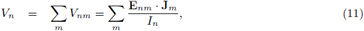

where **E**_*nm*_ is the electric field vector generated at brain location m when stimulating scalp electrode *n* with intensity *I_n_*. We can express (11) in matrix notation as:

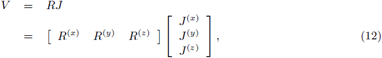

where

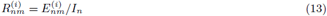

is the element at row *n* and column m of matrix *R^(i)^*.

Let us turn now to the stimulation case. We seek to write the net electric field generated in the brain when simultaneously stimulating at multiple scalp electrodes (indexed by *n*). Once again employing the superposition principle, we write the total electric field as the sum of the individual electric fields generated by stimulation at each electrode:

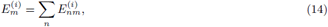

where 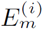 is the *i*th component of the net electric field at brain location *m*. From (13), we have that 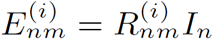, which we substitute into (14) to yield:

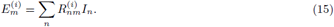

Writing (15) in matrix notation leads to:

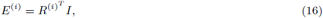

where *E^(i)^* is a vector whose mth element is 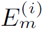, and *I* is a vector whose *n*th element is *I_n_*. From (2) and (16), we identify 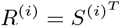, which is the desired result.

### Theorem 2.

*Reciprocal targeting. In a least-squares sense, the vector of scalp currents I^*^ which best recreates the neural current source distribution J is given by:*

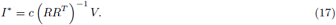

*Proof.* We seek to minimize the squared error between *E* and *cJ*, leading to the following optimization problem:

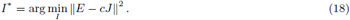

Let distance *D = ||E − cJ||^2^* denote the cost function to be minimized. Taking the derivative with respect to the vector of applied currents *I* leads to:

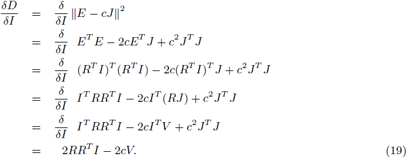

Setting (19) to zero leads to

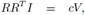

from which we arrive at the desired result (17).

## Reciprocal TES is equivalent to minimum-norm EEG source localization

Here we show that reciprocal TES targeting following (4) is actually the counterpart of the minimum-norm solution to the MEG/EEG source localization problem (Sarvas, 1987; Hämäläinen and Ilmoniemi, 1994). To see this, substitute (4) into (2), yielding the following expression for the electric field generated by reciprocal TES:

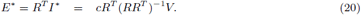

Consider now the inverse problem of finding the current density distribution *J* that gave rise to the observed pattern of scalp potentials *V = RJ*. This is an ill-posed problem without a unique solution. A common approach to solving such underdetermined systems is to identify the solution with minimum norm:

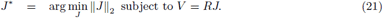

The solution to (21) is given by:

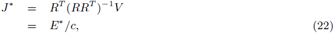

where the last line follows from (20).

## Algorithm for *L*^1^-constrained reciprocity

In order to identify the reciprocal TES montage that adheres to a hard constraint on the total current delivered to the head, we seek to solve the following constrained optimization problem:

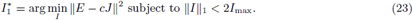

Following Tibshirani (1996), the non-differentiable *L*^1^ inequality constraint in (23) may be converted into a set of 2^*N*^ linear inequality constraints, where *N* is the number of scalp electrodes:

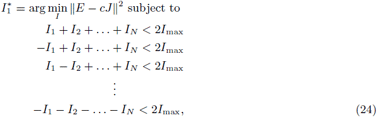

where the linear constraints span all possible “sign” combinations of the current vector *I*. In principle, the optimization problem (24) may then be solved using conventional quadratic programming. However, for many applications (including this one), the number of linear constraints 2^*N*^ is extremely high, making direct implementation of (24) intractable. To circumvent this, Tibshirani also proposed an iterative scheme to solving (24) that begins with the unconstrained solution *I^*^* and proceeds to update the solution, adding a single linear constraint at each iteration (Tibshirani, 1996). Empirical results (including the ones in this paper) show that convergence occurs long before all 2^*N*^ constraints have been added to the cumulative set of constraints. In general, the number of iterations required for the algorithm to converge is in the order of *N*. This efficient iterative scheme was employed here to solve for the *L*^1^-constrained reciprocal TES montage.

## Head model

The head model was constructed in the Brainstorm software package (Tadel et al., 2011) for Matlab (Mathworks, Natick, MA). The anatomy of the head was selected as the “ICBM-152” standard, which was constructed by non-linearly averaging 152 individual heads in a common coordinate system (Mazziotta et al., 1995; Fonov et al., 2011). The Brainstorm package relies on the FreeSurfer tool for segmenting the head into its constituent tissue categories (Fischl, 2012). The BEM was constructed with the following (default Brainstorm) parameters: skull: relative conductivity of 1 S/m, scalp: relative conductivity of 0.0125 S/m, brain: relative conductivity of 1 S/m. The model included 15,002 virtual sources (vertices) arranged along the pial surface. 64 scalp electrodes (63 free electrodes + 1 reference) were then placed on the scalp following the international 10/10 standard for EEG. To solve for the lead field from each source to each scalp electrode, Brainstorm employed the OpenMEEG tool (Gramfort et al., 2010). This yielded the 63-by-45006 matrix R, where the rows spanned electrodes and the columns spanned the 3 Cartesian orientations of each cortical source.

When simulating neural activations, the following model was used:

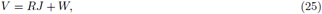

where *W* was an *N* − 1 = 63 element vector of spatially uncorrelated Gaussian noise added to the electrode array. All activated sources had a magnitude of 1 nA · m and an orientation perpendicular to the local cortical sheet. Unless otherwise stated, the standard deviation of *W* was selected to be 0.01 of the standard deviation of *RJ* (in other words, the nominal SNR was set to 100). It is well-known that in practice, the (single-trial) SNR of EEG data is quite low. Here, however, the SNR was deliberately increased in order to (i) confirm the theoretical findings under low-noise conditions, (ii) because, in practice, reciprocal TES is expected to be applied to either averaged or component data whose SNR is much higher than the single-trial SNR (see Discussion), and (iii) to determine the upper-bounds on performance of reciprocal TES. Neural activations consisted of a “seed” location and a fixed number of vertices *K* closest to the seed point. The seed location was initially selected by clicking on a vertex on the cortical surface. The value of *K* was selected (see below for specific values) such that the resulting activation spanned the desired area of cortex.

Unless otherwise indicated, the value of parameter c was set to 10^10^. All TES montages were restricted to a total current delivered of 2 mA. This was achieved using Eq. (7) for the “optimal” (unconstrained) and “naive” reciprocity, and using Eq. (8) for *L*^1^-constrained reciprocity.

To measure focality, we estimated the radial distance from the target at which the total electric field contained within that radius dropped to half of the total electric field (across all locations). When estimating the number of electrodes recruited by reciprocal TES, we computed the proportion of electrodes carrying at least 5% of the total applied current.

*Figure 1*. The seed point was located at (0.038,−0.038,0.11), and *K* = 25.

*Figure 2*. There were two seed points: (−0.028,0.0083,0.12) and (−0.025,−0.012,0.13), each with *K* = 25.

*Figure 3*. For panels A-D, the seed location and *K* values were selected as follows: left superior temporal gyrus (STG) (0.013,0.067,0.049), *K* = 20; left dorsolateral prefrontal cortex (DLPFC) (0.076,0.040,0.072), *K* = 20; left superior parietal lobule: (−0.037,0.017,0.12), *K* = 15; left middle temporal area (MT): (-0.064,0.044,0.056), *K* = 15. For the example activation of visual cortex (Panels E-G), the two seed points (each with *K* = 25) were located at (−0.070, −0.0084, 0.069) and (−0.070,−0.000018,0.066). When simulating activations from all 15,002 sources (Panel H), we iterated over all vertices, each time computing the *K* = 50 vertices closest to the seed vertex. On average, the radius of the minimum sphere which bounded the activated vertices was 1.2 cm.

*Figure 4*. Please refer to the values from Figure 3, Panel H above.

*Figure 5*. The seed point was located at (0.017, 0.041, 0.11), and *K* = 20. The orientation of the tangential source was set to (0,1, 0).

## Public availability of data

The head model, as well as the code to perform reciprocal TES, are available for download in Matlab format at: TBD.

## References

Antal, A. and Paulus, W. (2013). Transcranial alternating current stimulation (tacs). Frontiers in human neuroscience, 7.

Berényi, A., Belluscio, M., Mao, D., and Buzsáki, G. (2012). Closed-loop control of epilepsy by transcranial electrical stimulation. Science, 337(6095):735–737.

Bergmann, T. O., Karabanov, A., Hartwigsen, G., Thielscher, A., and Siebner, H. R. (2016). Combining non-invasive transcranial brain stimulation with neuroimaging and electrophysiology: Current approaches and future perspectives. NeuroImage.

Bestmann, S. and Feredoes, E. (2013). Combined neurostimulation and neuroimaging in cognitive neuroscience: past, present, and future. Annals of the New York Academy of Sciences, 1296(1):11–30.

Bikson, M., Grossman, P., Thomas, C., Zannou, A. L., Jiang, J., Adnan, T., Mourdoukoutas, A. P., Kronberg, G., Truong, D., Boggio, P., et al. (2016). Safety of transcranial direct current stimulation: evidence based update 2016. Brain Stimulation, 9(5):641–661.

Bikson, M., Inoue, M., Akiyama, H., Deans, J. K., Fox, J. E., Miyakawa, H., and Jefferys, J. G. (2004). Effects of uniform extracellular dc electric fields on excitability in rat hippocampal slices in vitro. The Journal of physiology, 557(1):175–190.

Brunoni, A. R., Nitsche, M. A., Bolognini, N., Bikson, M., Wagner, T., Merabet, L., Edwards, D. J., Valero-Cabre, A., Rotenberg, A., Pascual-Leone, A., et al. (2012). Clinical research with transcranial direct current stimulation (tdcs): challenges and future directions. Brain stimulation, 5(3):175–195.

Cancelli, A., Cottone, C., Tecchio, F., Truong, D. Q., Dmochowski, J., and Bikson, M. (2016). A simple method for eeg guided transcranial electrical stimulation without models. Journal of neural engineering, 13(3):036022.

Delorme, A. and Makeig, S. (2004). Eeglab: an open source toolbox for analysis of single-trial eeg dynamics including independent component analysis. Journal of neuroscience methods, 134(1):9–21.

Dmochowski, J. P., Datta, A., Bikson, M., Su, Y., and Parra, L. C. (2011). Optimized multi-electrode stimulation increases focality and intensity at target. Journal of neural engineering, 8(4):046011.

Dmochowski, J. P., Sajda, P., Dias, J., and Parra, L. C. (2012). Correlated components of ongoing eeg point to emotionally laden attention-a possible marker of engagement? Frontiers in human neuroscience, 6(112).

Dostrovsky, J., Levy, R., Wu, J., Hutchison, W., Tasker, R., and Lozano, A. (2000). Microstimulation-induced inhibition of neuronal firing in human globus pallidus. Journal of neurophysiology, 84(1):570–574.

Faria, P., Fregni, F., Sebastião, F., Dias, A. I., and Leal, A. (2012). Feasibility of focal transcranial dc polarization with simultaneous eeg recording: preliminary assessment in healthy subjects and human epilepsy. Epilepsy & Behavior, 25(3):417–425.

Fernández-Corazza, M., Turovets, S., Luu, P., Anderson, E., and Tucker, D. (2016). Transcranial electrical neuromodulation based on the reciprocity principle. front. Psychiatry, 7:87.

Fischl, B. (2012). Freesurfer. Neuroimage, 62(2):774–781.

Fonov, V., Evans, A. C., Botteron, K., Almli, C. R., McKinstry, R. C., Collins, D. L., Group, B. D. C., et al. (2011). Unbiased average age-appropriate atlases for pediatric studies. NeuroImage, 54(1):313–327.

Gramfort, A., Luessi, M., Larson, E., Engemann, D. A., Strohmeier, D., Brodbeck, C., Parkkonen, L., and Hämäläinen, M. S. (2014). Mne software for processing meg and eeg data. Neuroimage, 86:446–460.

Gramfort, A., Papadopoulo, T., Olivi, E., and Clerc, M. (2010). Openmeeg: opensource software for quasistatic bioelectromagnetics. Biomedical engineering online, 9(1):1.

Guggenmos, D. J., Azin, M., Barbay, S., Mahnken, J. D., Dunham, C., Mohseni, P., and Nudo, R. J. (2013). Restoration of function after brain damage using a neural prosthesis. Proceedings of the National Academy of Sciences, 110(52):21177–21182.

Hallez, H., Vanrumste, B., Grech, R., Muscat, J., De Clercq, W., Vergult, A., D’Asseler, Y., Camilleri, K. P., Fabri, S. G., Van Huffel, S., et al. (2007). Review on solving the forward problem in eeg source analysis. Journal of neuroengineering and rehabilitation, 4(1):1.

Hämäläinen, M. S. and Ilmoniemi, R. J. (1994). Interpreting magnetic fields of the brain: minimum norm estimates. Medical & biological engineering & computing, 32(1):35–42.

Haufe, S., Meinecke, F., Görgen, K., Dähne, S., Haynes, J.-D., Blankertz, B., and Bießmann, F. (2014). On the interpretation of weight vectors of linear models in multivariate neuroimaging. Neuroimage, 87:96–110.

Helmholtz, H. v. (1853). Ueber einige gesetze der vertheilung elektrischer straöme in kaörperlichen leitern, mit anwendung auf die thierisch-elektrischen versuche (schluss.). Annalen der Physik, 165(7):353–377.

Huang, Y., Parra, L. C., and Haufe, S. (2015). The new york head−a precise standardized volume conductor model for eeg source localization and tes targeting. NeuroImage.

Jackson, J. D. (1999). Classical electrodynamics. Wiley.

Jimbo, Y., Kasai, N., Torimitsu, K., Tateno, T., and Robinson, H. P. (2003). A system for mea-based multisite stimulation. Biomedical Engineering, IEEE Transactions on, 50(2):241–248.

Jung, Y.-J., Kim, J.-H., and Im, C.-H. (2013). Comets: a matlab toolbox for simulating local electric fields generated by transcranial direct current stimulation (tdcs). Biomedical engineering letters, 3(1):39–46.

Kent, A. R. and Grill, W. M. (2013). Neural origin of evoked potentials during thalamic deep brain stimulation. Journal of neurophysiology, 110(4):826–843.

Lempka, S. F. and McIntyre, C. C. (2013). Theoretical analysis of the local field potential in deep brain stimulation applications. PloS one, 8(3):e59839.

Maynard, E. M., Nordhausen, C. T., and Normann, R. A. (1997). The utah intracortical electrode array: a recording structure for potential brain-computer interfaces. Electroencephalography and clinical neurophysiology, 102(3):228–239.

Mazziotta, J. C., Toga, A. W., Evans, A., Fox, P., and Lancaster, J. (1995). A probabilistic atlas of the human brain: Theory and rationale for its development: The international consortium for brain mapping (icbm). Neuroimage, 2(2):89–101.

Nunez, P. L. and Srinivasan, R. (2006). Electric fields of the brain: the neurophysics of EEG. Oxford University Press, USA.

Opitz, A., Falchier, A., Yan, C.-G., Yeagle, E., Linn, G., Megevand, P., Thielscher, A., Milham, M., Mehta, A., and Schroeder, C. (2016). Spatiotemporal structure of intracranial electric fields induced by transcranial electric stimulation in human and nonhuman primates. bioRxiv, page 053892.

Osorio, I., Frei, M. G., Manly, B. F., Sunderam, S., Bhavaraju, N. C., and Wilkinson, S. B. (2001). An introduction to contingent (closed-loop) brain electrical stimulation for seizure blockage, to ultrashort-term clinical trials, and to multidimensional statistical analysis of therapeutic efficacy. Journal of Clinical Neurophysiology, 18(6):533–544.

Parra, L. C., Spence, C. D., Gerson, A. D., and Sajda, P. (2005). Recipes for the linear analysis of eeg. Neuroimage, 28(2):326–341.

Pascual-Marqui, R. D. (1999). Review of methods for solving the eeg inverse problem. International journal of bioelectromagnetism, 1(1):75–86.

Radman, T., Ramos, R. L., Brumberg, J. C., and Bikson, M. (2009). Role of cortical cell type and morphology in subthreshold and suprathreshold uniform electric field stimulation in vitro. Brain stimulation, 2(4):215–228.

Rahman, A., Reato, D., Arlotti, M., Gasca, F., Datta, A., Parra, L. C., and Bikson, M. (2013). Cellular effects of acute direct current stimulation: somatic and synaptic terminal effects. The Journal of physiology, 591(10):2563–2578.

Rosenberg, D., Mauguiere, F., Catenoix, H., Faillenot, I., and Magnin, M. (2009). Reciprocal thalamocortical connectivity of the medial pulvinar: a depth stimulation and evoked potential study in human brain. Cerebral Cortex, 19(6):1462–1473.

Rosin, B., Slovik, M., Mitelman, R., Rivlin-Etzion, M., Haber, S. N., Israel, Z., Vaadia, E., and Bergman, H. (2011). Closed-loop deep brain stimulation is superior in ameliorating parkinsonism. Neuron, 72(2):370–384.

Rush, S. and Driscoll, D. A. (1969). Eeg electrode sensitivity-an application of reciprocity. Biomedical Engineering, IEEE Transactions on, (1):15–22.

Sarvas, J. (1987). Basic mathematical and electromagnetic concepts of the biomagnetic inverse problem. Physics in medicine and biology, 32(1):11.

Siebner, H. R., Bergmann, T. O., Bestmann, S., Massimini, M., Johansen-Berg, H., Mochizuki, H., Bohning, D. E., Boorman, E. D., Groppa, S., Miniussi, C., et al. (2009). Consensus paper: combining transcranial stimulation with neuroimaging. Brain, stimulation, 2(2):58–80.

Silvanto, J., Muggleton, N., and Walsh, V. (2008). State-dependency in brain stimulation studies of perception and cognition. Trends in cognitive sciences, 12(12):447–454.

Tadel, F., Baillet, S., Mosher, J. C., Pantazis, D., and Leahy, R. M. (2011). Brainstorm: a user-friendly application for meg/eeg analysis. Computational intelligence and neuroscience, 2011:8.

Thielscher, A., Antunes, A., and Saturnino, G. B. (2015). Field modeling for transcranial magnetic stimulation: A useful tool to understand the physiological effects of tms? In 2015 37th Annual International Conference of the IEEE Engineering in Medicine and Biology Society (EMBC), pages 222–225. IEEE.

Thut, G., Ives, J. R., Kampmann, F., Pastor, M. A., and Pascual-Leone, A. (2005). A new device and protocol for combining tms and online recordings of eeg and evoked potentials. Journal of neuroscience methods, 141(2):207–217.

Tibshirani, R. (1996). Regression shrinkage and selection via the lasso. Journal of the Royal Statistical Society. Series B (Methodological), pages 267–288.

Trebuchon, A., Guye, M., Tcherniack, V., Tramoni, E., Bruder, N., and Metellus, P. (2012). [interest of eeg recording during direct electrical stimulation for brain mapping function in surgery]. In Annales francaises d’anesthesie et de reanimation, volume 31, pages e87–90.

Truong, D. Q., Hüber, M., Xie, X., Datta, A., Rahman, A., Parra, L. C., Dmochowski, J. P., and Bikson, M. (2014). Clinician accessible tools for gui computational models of transcranial electrical stimulation: Bonsai and spheres. Brain stimulation, 7(4):521–524.

Uhlhaas, P. J. and Singer, W. (2006). Neural synchrony in brain disorders: relevance for cognitive dysfunctions and pathophysiology. Neuron, 52(1):155–168.

Uhlhaas, P. J. and Singer, W. (2012). Neuronal dynamics and neuropsychiatric disorders: toward a translational paradigm for dysfunctional large-scale networks. Neuron, 75(6):963–980.

Wagner, S., Lucka, F., Vorwerk, J., Herrmann, C., Nolte, G., Burger, M., and Wolters, C. H. (2016). Using reciprocity for relating the simulation of transcranial current stimulation to the eeg forward problem. NeuroImage.

Zaehle, T., Rach, S., and Herrmann, C. S. (2010). Transcranial alternating current stimulation enhances individual alpha activity in human eeg. PloS one, 5(11):e13766.

